# Shaping and probing living tissues with magnetic bioprinting

**DOI:** 10.1101/2025.09.18.677128

**Authors:** Noam Demri, Polina Petrova Tsvetkova, Carine Vias, Giacomo Gropplero, Simon Dumas, Fanny Cayrac, Stéphanie Descroix, Claire Wilhelm

## Abstract

Mechanical and geometric cues play a crucial role *in vivo*, regulating both morphogenetic processes and proper tissue function. This is particularly evident in skeletal muscle, where aligned architecture is essential for myogenesis and functional force generation. However, precisely engineering tissue geometry at both macroscopic and microscopic scales while simultaneously controlling internal mechanical forces remains a significant challenge. In this study, we introduce a magnetic tissue engineering platform based on a magnetic bioprinting technique, enabling precise control of biophysical cues that guide *in vitro* tissue morphogenesis. Applied here to skeletal muscle, this approach allows for the rapid fabrication of cohesive tissues in any desired shape using cells labeled with magnetic nanoparticles. Additionally, multiple cell types can be incorporated and spatially organized within the same construct through magnetic segregation. Optimizing tissue geometry further enables magnetic actuation, including the ability to trap and maintain tissue shape over time. Furthermore, this magnetic platform facilitates the investigation of how tissue architecture influences mechanical properties, such as resistance to rupture. Overall, this study highlights the significant potential of magnetic bioprinting and stimulation for controlling tissue morphology and advancing biomechanical research.

## I. Introduction

As soon as embryogenesis begins, cells are subjected to mechanical and geometrical constraints that guide morphogenesis [1,2]. Coupled with biochemical signals, these structural cues diverge locally, directing cells to differentiate in a spatially dependent manner and ultimately shaping the different organs of the body [3]. Understanding and replicating these morphogenic processes has become essential for developing systems that accurately recreate *in vitro* the complex *in vivo* tissue organization. Advances in regenerative medicine have motivated efforts to engineer physiologically relevant tissues at large scale [3], while microphysiological systems [4–13], such as organs-on-chips, have successfully reproduced key aspects of tissue microenvironments. However, achieving full control over a tissue’s macroscopic shape, internal cellular organization, and mechanical forces remains challenging, particularly at larger scales[14–16].

Several approaches have been developed to impose tissue geometry, including matrix moulds, to shape cells or tissues [12,13,17–19], and bioprinting, where cells are seeded into a biocompatible scaffold that is either extruded [20–22] or photopolymerized [23,24] in a controlled geometry. These platforms enable the fabrication of complex tissue architectures, sometimes even accurately replicating the overall geometry of entire tissues and even organs [25]. Despite these advancements, scaffold-based approaches have limitations. They can restrict cell density [26], and while the initial geometry is precisely controlled, the dynamic evolution of the tissue’s internal cellular architecture and stresses remains challenging to regulate. These structural and mechanical cues are essential to morphogenesis. Skeletal muscle tissue provides a clear example, as alignment is present at every scale - from the entire organ spanning from tendon to tendon to actin-myosin complexes within multinucleated myofibers, all oriented in the direction of muscle contraction[27,28]. This uniform alignment optimizes force generation and enables movement. Moreover, aligned geometry and mechanical forces are not only functionally coupled, but are also essential in myogenesis, where mononucleated myoblasts fuse together to form aligned myotubes. Such coupling of structural and biomechanical cues plays a crucial role in many other tissues and physiological processes. This includes the development and function of smooth [29] and cardiac muscles but extends well beyond muscle tissue. For instance, endothelial cells in tubular tissues display a specific apical-basal polarity and respond to fluid shear stresses[8]. Similarly, gut organoid development, particularly the formation of crypt-villus structures, has been shown to improve under cyclic mechanical stretching[30]. These examples underscore the widespread significance of mechanical and geometric cues in tissue formation and function.

Replicating *in vivo* dynamic mechanical stresses has therefore become a key challenge in tissue engineering. Different strategies have been developed to mimic the physiological conditions, including pressure-controlled flows in microfluidic devices [8,18], or engineered systems to constrain, compress or stretch [31–37] tissues.

A promising strategy for controlling both structural and mechanical cues in tissue engineering involves the use of magnetic forces. Iron oxide nanoparticles, widely used in imaging (e.g. as MRI contrast agents) and therapeutic (e.g. for drug delivery or hyperthermia treatment) applications[38–40], can be efficiently internalized by various cell types via the endocytic pathway [31,41], without impairing cellular functions. Once internalized, these nanoparticles enable remote cellular manipulation through external magnetic fields, providing a non-invasive means of exerting precise mechanical forces on cells [42–44]. Building on this principle, magnetic tissue engineering approaches have emerged, beginning with the development of magnetically produced cell sheets [45]. This technology was subsequently adapted to generate simple tissue geometries, such as rods, spheroids or rings [31,46–48], with the added potential for remote magnetic manipulation [31,36,48–51]. However, the full potential of magnetic tissue engineering remains untapped, particularly in achieving more complex tissue architectures and in precise dynamic biomechanical control.

To address these challenges, we developed an innovative magnetic tissue engineering platform that enables precise, remote control over both tissue geometry and biomechanics. A key feature of this platform is a magnetic bioprinting approach capable of generating tissues of arbitrary shapes while maintaining control at both macroscopic and microscopic levels. We demonstrate that optimizing tissue geometry enhances the potential for further magnetic actuation, enabling shape retention over time. Using skeletal muscle as a proof of concept, we show that this platform allows for the controlled evolution of tissue shape, as well as the internal organization of cells, including their orientation and spatial positioning via magnetic segregation. Additionally, this system allows for the assessment of tissue mechanical properties, specifically resistance to fracture, which is found to be dependent on the variations of stimulation parameters, including the speed, amplitude, and duration of the applied stress, as well as the cell type used.

## II. Results

### Versatile magnetic bioprinting for custom tissue architecture

Magnetic tissue engineering, particularly magnetic bioprinting, requires both magnetically labeled cells and precisely controlled magnetic fields to organize them. To achieve the magnetic labelling, C2C12 mouse myoblasts cells were exposed to superparamagnetic iron oxide nanoparticles at a 2 mM iron concentration for 30 minutes, once daily, over the three days prior to printing. This labeling procedure, referred to as NP3, allows cells to internalize approximately 20 pg of iron per cell via the endocytic pathway, as previously described [31], therefore enabling their remote actuation through magnetic forces. To precisely control the magnetic field distribution, NiFe micropatterns with the desired geometry were fabricated using photolithography followed by NiFe electrodeposition onto a copper-coated glass slide (Fig. 1A). NiFe patterns grow onto the exposed copper regions defined by photolithography while the remaining substrate is protected by a resin layer, forming 50 µm-thick patterns that replicate the exact same shape of the photomask design [52]. Next, to perform magnetic bioprinting, a suspension of magnetically labeled cells is deposited in a well with a thin coverslip at the bottom, which is placed onto these micropatterns. The system is exposed to a strong magnetic field generated by two opposing magnets positioned to minimize field gradient, as done previously [53]. While the NiFe patterns are not magnetic on their own, they act as magnetic concentrators when exposed to the external magnetic field, locally amplifying magnetic forces and directing the cells towards them [31]. After a 3h incubation period and subsequent removal from the magnetic field, the cell aggregates self-organize into cohesive tissues that replicate the pattern geometry (Fig. 1B). This was demonstrated using patterns of various shapes, including crosses (Fig. 1C), stars (Fig. 1D), triangles (Fig. 1E) or droplets (Fig. 1F). More complex structures, such as wrench-shaped geometries, could also be fabricated, with the ability to modulate key geometrical features of the printed tissues, such as width (Fig. 1G), length (Fig. 1H), and the shape of the clamps at each extremity (Fig. 1I). These results highlight the versatility of the magnetic bioprinting technique for engineering tissues with customizable shapes and dimensions.

**Figure 1.**
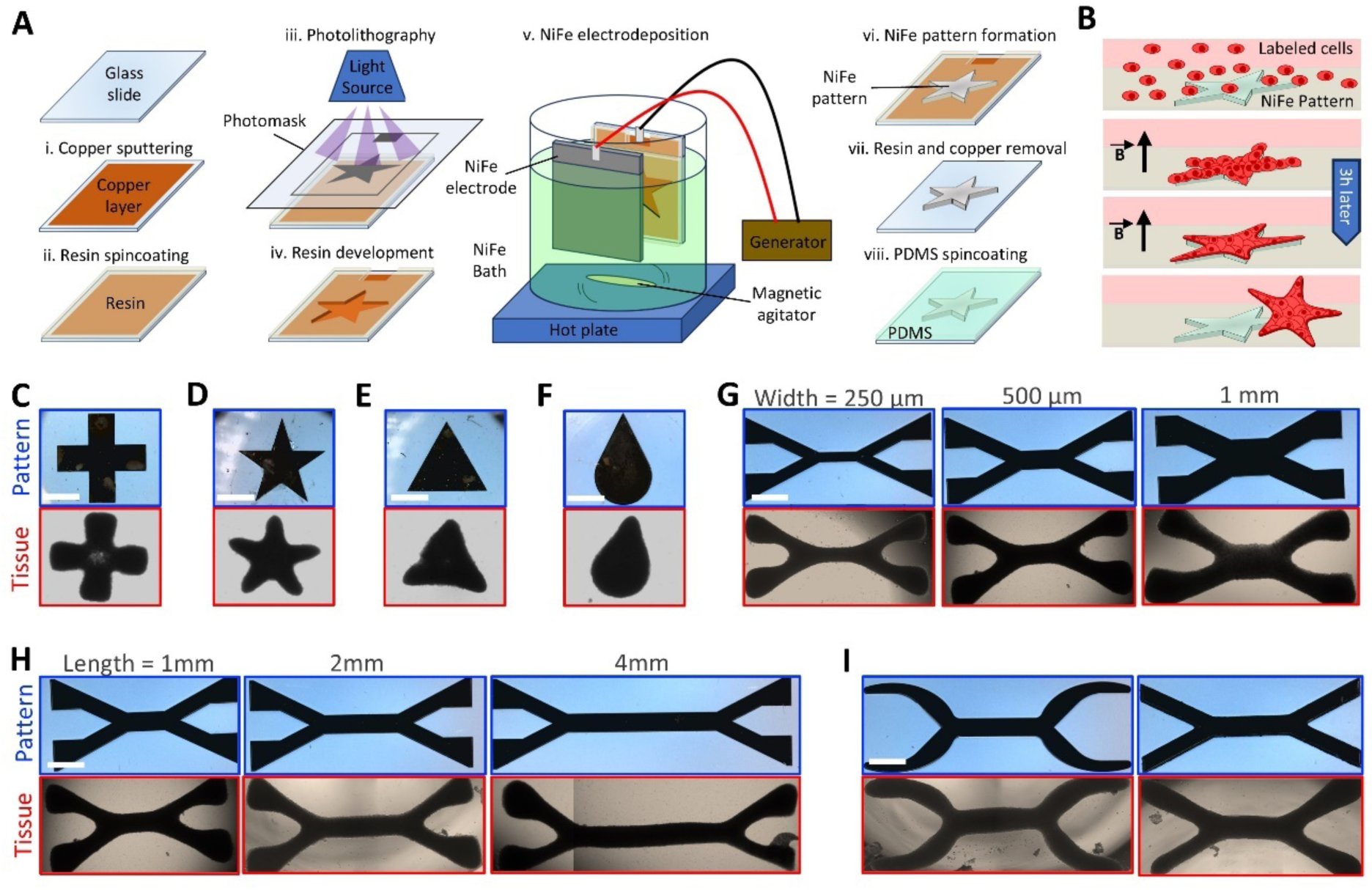
Magnetic bioprinting of cellular tissues with customizable geometries. (A-B) Schematics of (A) the NiFe pattern fabrication process and (B) the magnetic bioprinting process. (C-F) Magnetic patterns (top) and the resulting printed tissues made of C2C12 cells (bottom) in the shapes of (C) a cross, (D) a star, (E) a triangle, and (F) a droplet. (G-I) Wrench-shaped magnetic patterns (top) and the resulting printed tissues (bottom) demonstrating variations in central fiber (G) width, (H) length, and (I) clamp shape. (Scale bar = 1mm).

### Tissue maturation drives contraction towards spherical geometry

Once the magnetic field is removed, the printed tissues readily detach from the coverslip at the bottom of the well, which was coated with an anti-adherence solution before bioprinting. The tissues can therefore easily move away from the position of the patterns below the coverslip, which facilitates their collection and manipulation. However, it also means they are not constrained anymore. While magnetic bioprinting ensures precise control over tissue shape at the time of printing (day 0), maintaining this shape over time proved significantly more challenging, especially for contractile cells such as myoblasts. When left free-floating in a non-adherent dish, the tissues gradually transition into a spherical shape, likely due to the overall tissue surface tension and cellular contractility (Fig. 2A, Supplementary Fig. 1A, Supplementary Movies 1 and 2). Since preserving tissue geometry over time is crucial, this “spherification” process needed to be investigated, and strategies to prevent it had to be developed. To address this, we focused on cross-shaped tissues, as their geometry is very distinct from that of a circle and can be easily modulated. The first strategy to mitigate shape loss was to vary tissue scale. Cross-shaped NiFe patterns were made with branch lengths of 1, 2 and 4 mm, all 500 µm wide. While the magnetic bioprinting process successfully generated these larger centimeter-scale tissues, they invariably transitioned towards a circular shape (Fig. 2A, Supplementary Fig. 1A). Smaller crosses compacted by retracting their branches toward the center, whereas larger ones folded onto themselves before rounding up. Quantification of the projected tissue area over six days (Fig. 2B) revealed an exponential decay, with a time constant of approximately 1.5 days across all cross sizes. Further analysis of tissue circularity (Fig. 2C) and solidity (Fig. 2D) confirmed a progressive evolution toward a convex sphere at the same time as they become more and more compact.

**Figure 2.**
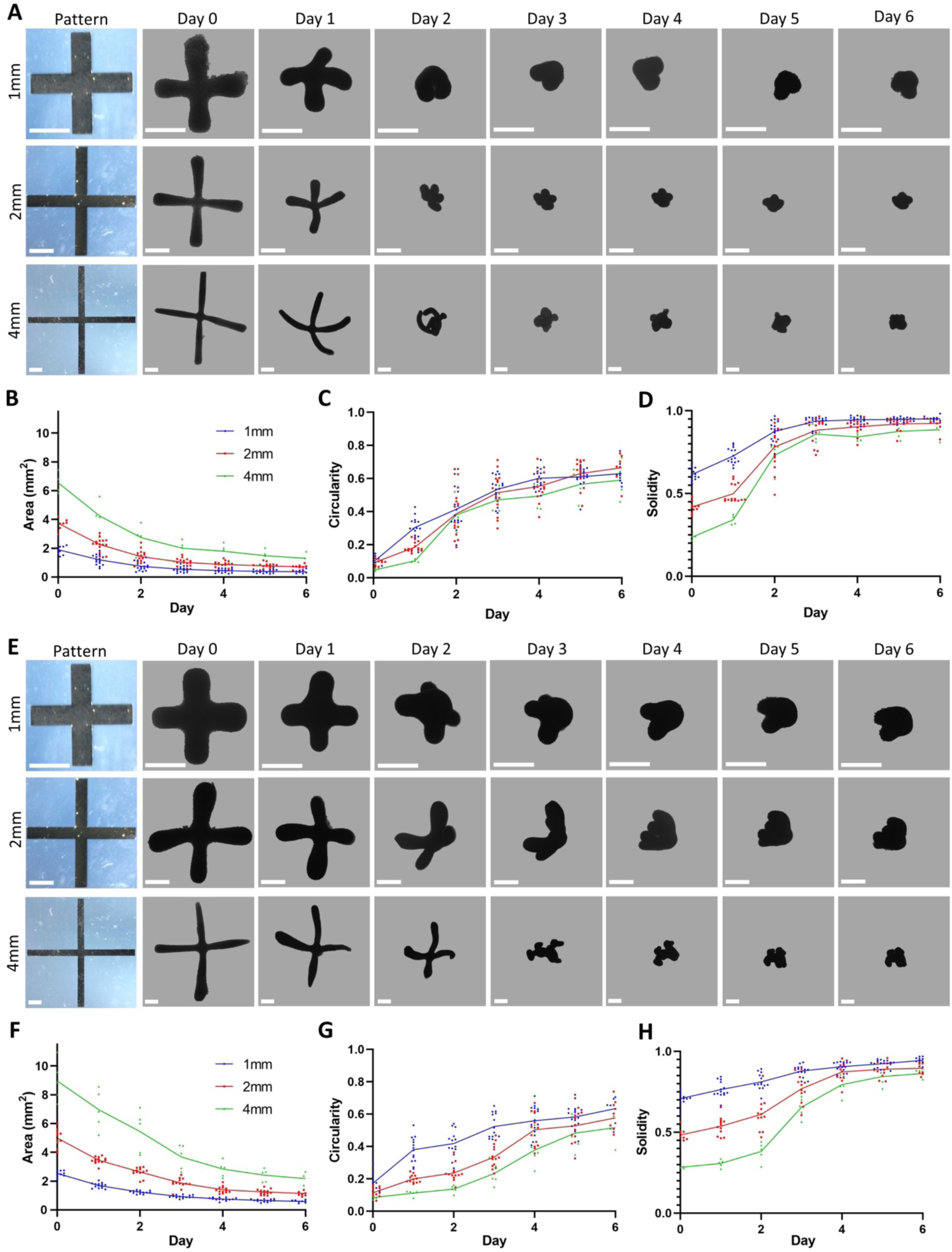
Cross-shaped tissues naturally mature toward a spherical shape. (A) Brightfield images from day 0 to day 6 of free-floating tissues printed using cross patterns with branch lengths of 1, 2 and 4 mm, at a cell density of 100,000 cells per mm² of pattern. Evolution of their (B) area, (C) circularity and (D) solidity over time. (E) Brightfield images from day 0 to day 6 of free-floating tissues printed using the same cross patterns but with a higher cell density of 200,000 cells per mm² of pattern. Evolution of their (F) area, (G) circularity and (H) solidity over time. Images of the tissues were segmented to remove the background. (Scale bar = 1 mm).

As an alternative strategy to better maintain their initial shape, we increased tissue thickness rather than length. To achieve this, the tissues that were previously printed at a cell density of 100,000 cells per mm^2^ of NiFe pattern were now printed at twice this density, resulting in thicker tissues (Fig. 2E, Supplementary Fig. 1B). While the thickness increase extended the time constant to up to three 3 days, the area still shrank exponentially (Fig. 2F) with these tissues eventually reaching the same levels of circularity (Fig. 2G) and convexity (Fig. 2H) of the thinner tissues.

Since modifying tissue geometry only appeared to slow down, rather than prevent shape transformation, other strategies to preserve the 3D architecture were explored - specifically, maintaining the shape by applying the magnetic force continuously during maturation or embedding the tissue within a matrix. In the initial bioprinting process, tissues were collected after three hours of exposure to the magnetic field, forming a structure that precisely matched the underlying NiFe patterns (Fig. 3A-B). Given that the pattern-mediated magnetic forces drive cell assembly, an initial hypothesis was that maintaining the tissues under these forces over longer periods of time might help to preserve their shapes. However, when tissues remained on the patterns placed within the magnets for six days, they still lost their original geometry, eventually compacting into spherical aggregates on top of the patterns (Fig. 3C-D). This suggests that the contractile forces driving shape transition are stronger than pattern-induced magnetic forces, previously evaluated at approximately 1nN on a single cell [31,52,53].

**Figure 3.**
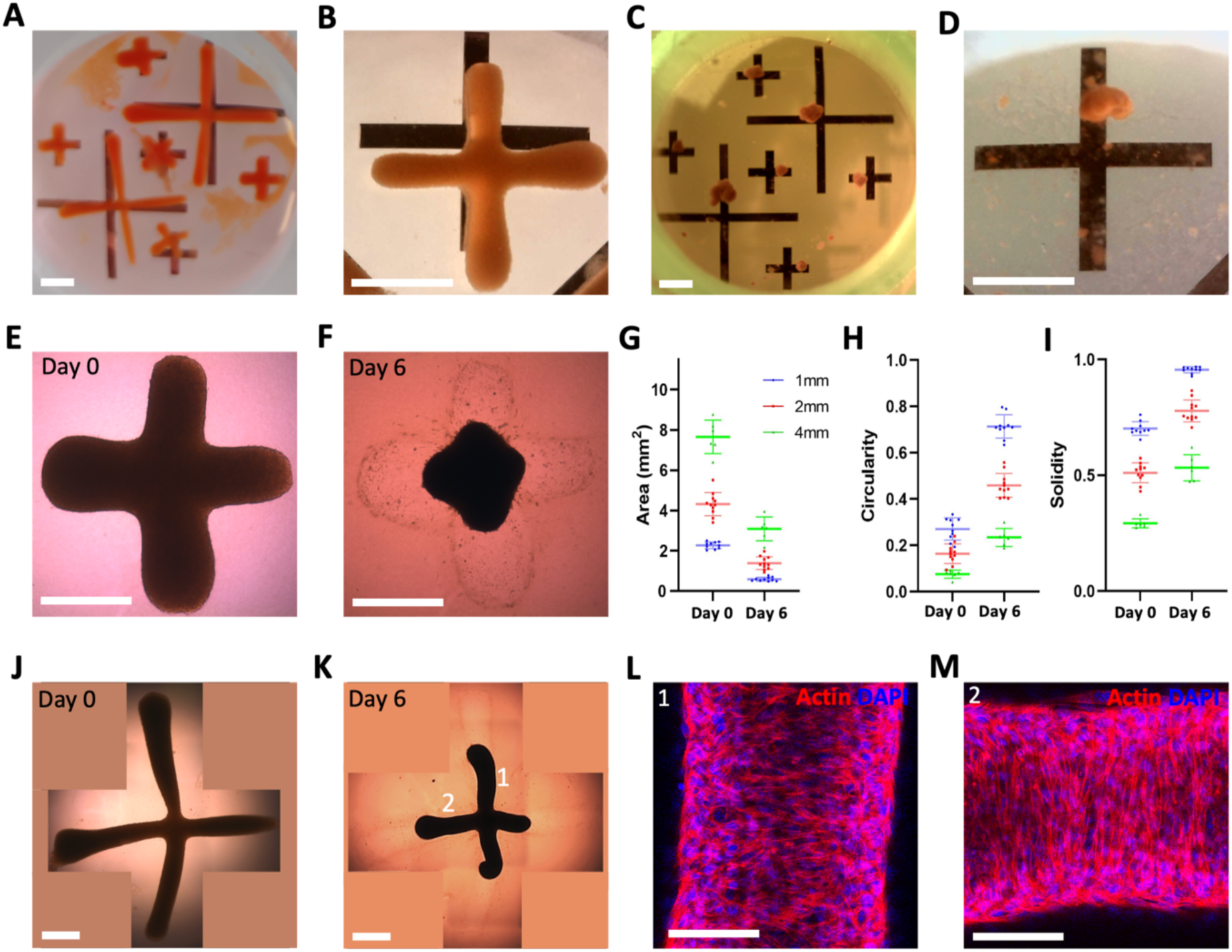
Tissue shape cannot be fully preserved by magnetic patterns or a collagen matrix. (A-B) Tissues remaining on top the patterns in a bioprinting well after 3 hours and (C-D) 6 days between magnets, shown at both whole-well scale and at the level of a cross with 2 mm branches. (Scale bar = 2 mm) (E-F) Brightfield images of a tissue made with cross patterns with 1 mm branches and 200,000 cells per mm^2^ of pattern, trapped in a collagen matrix, on (E) day 0 and (F) day 6. (G) Area, (H) circularity, (I) solidity of tissues made with 1-, 2- and 4-mm long branches and 200,000 cells per mm^2^ of pattern on days 0 and 6. Brightfield of a tissue made with cross patterns with 4 mm branches and 200,000 cells per mm^2^ of pattern, trapped in a collagen matrix, on (J) day 0 and (K) day 6. (Scale bar = 1 mm) Tissues were not segmented to show the deformation of the matrix. As tissues were reconstructed from multiple frames, blank parts were filled with a uniform background color. (L-M) Confocal images of slices of two perpendicular branches of a cross-shaped tissue trapped in collagen on day 6, with nuclei in blue and actin in red. (Scale bar = 100 µm)

A distinctive feature of magnetic bioprinting is its scaffold-free nature, setting it apart from most other bioprinting technique [20–23]. While the absence of supportive matrix allows for greater flexibility in tissue assembly, it also means that tissues lack external structural reinforcement. We therefore investigated whether embedding printed tissues in a hydrogel matrix could help preserve their shape. To test this, tissues were bioprinted as usual, but instead of being cultured free-floating in medium, they were embedded in a 2 mg.mL^-1^ collagen matrix (Fig. 3E). After 24 hours, cells at the tissue periphery began invading the gel (Supplementary Fig. 2), but the main structure remained cohesive and simply detached from the gel, leaving an imprint of its original shape within the collagen matrix on day 6 (Fig. 3F). While embedding in collagen modestly slowed shape transformation for small cross-shaped tissues (area, Fig. 3G; circularity, Fig. 3H; and solidity, Fig. 3I), larger ones retained their shape better than those observed in free-floating tissues (Fig. 3J-K). Although they also detached from the matrix and compacted, the matrix prevented the large structures from folding onto themselves. However, despite this macroscopic improvement, confocal imaging revealed a misalignment between microscopic cellular organization and tissue geometry. Instead of aligning along the branches of the cross, myoblasts oriented perpendicular to them (Fig. 3L-M). Since myoblast alignment is crucial for proper differentiation and fusion, and usually matches the alignment of the macroscopic geometry *in vivo,* this discrepancy suggests that the matrix alone is insufficient to fully preserve and control both the shape and internal architecture of magnetically printed tissues.

### Magnetic trapping stabilizes tissue architecture

While the magnetic forces exerted by the patterns were insufficient to balance the contraction forces, properly focused magnetic forces could still serve to mechanically constrain the tissues. To achieve this, cross-shaped patterns were redesigned to include clamps at the end of each branch to engineer clip-on cross-shaped tissues (Fig. 4A and Supplementary Fig. 3), enabling magnetic entrapment by gripping onto four magnetized needles. Chips made of Ecoflex were specifically designed for that effect, featuring needles at 90° angles and directed toward the chip’s center. Each needle was magnetized by an adjacent external magnet (Fig. 4B). Simulations indicated that the specific arrangement of the external magnets had little effect on the magnitude of the magnetic field gradient near the needles, which remained of about 10^4^ T/m (Fig. 4C and Supplementary Fig. 4). This resulted in forces of approximately 10 nN on NP3-labeled cells in the vicinity of the needles, whereas forces would progressively tend to 0 at the center of the chip. However, the magnet configuration influences the orientation of the magnetic field. An alternating arrangement of north and south poles generated a cross-like pattern between the needles. This configuration was chosen based on the expectation that magnetically printed tissues would align along the field lines. This was verified empirically: cross-shaped tissues transferred into the chip immediately reoriented themselves and clipped onto the needles (Fig. 4D). Furthermore, the collagen coating applied to the needles facilitated tissue adhesion. After overnight incubation, the tissues remained securely attached to the needles even after the external magnets were removed. This approach effectively constrained tissue movement while preserving the initial printed geometry.

**Figure 4.**
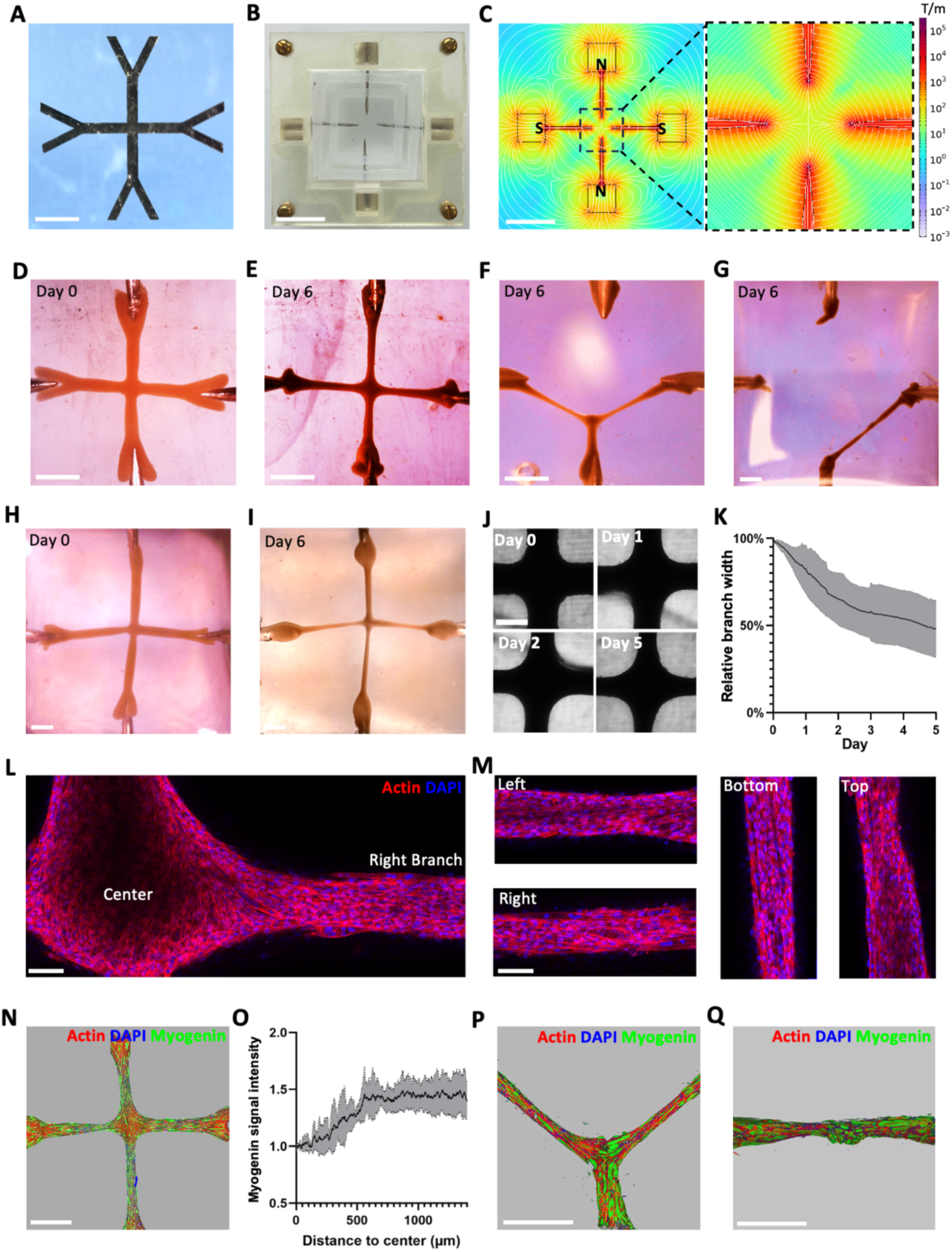
Clip-on cross-shaped tissues maintain their shape over time. (A) Clip-on cross-shaped pattern. (Scale bar = 2 mm). (B) Custom-designed chip for trapping the tissue, with (C) the corresponding magnetic field gradient and lines when magnets polarities (S for South and N for North) were alternated. (Scale bar = 1 cm). (D-E) Magnetically trapped tissue generated from clip-on cross patterns with 2 mm-long branches and 200,000 cells per mm^2^ of pattern, shown on (D) day 0 and (E) day 6. (F-G) Examples of broken magnetically trapped tissues produced from clip-on cross patterns using a lower density of 100,000 cells per mm^2^ of pattern, with (F) 2 mm and (G) 4 mm-long branches. (H-I) Magnetically trapped tissue formed from clip-on cross patterns with 4 mm-long branches and 200,000 cells per mm^2^ of pattern, with reinforced clamps, shown at (H) day 0 and (I) day 6 (Scale bar = 2 mm). (J-K) Evolution of branch width in cross-shaped tissues from day 0 to day 5, shown in (J) brightfield microscopy images and (K) plotted relative to the initial width (n=3) (Scale bar = 500 µm). (L-M) Confocal imaging of actin filaments in (L) the central region and (M) the 2 mm-long branches of a clip-on cross-shaped tissue on day 6. (Scale bar = 100 µm) (N-O) Spatial distribution of myogenin within a cross-shaped tissue with 2 mm-long branches on day 6, visualized in (N) a confocal 3D reconstruction and (O) plotted after normalization as a function of the distance from the center (n=5). (P-Q) 3D confocal reconstructions of 6-day-old trapped cross-shaped tissues where (P) one or (Q) two branches had broken. Actin is represented in red, nuclei in blue, and myogenin in green. (Scale bar = 1 mm)

As the tissues were cultured within the chips in differentiation medium, some successfully retained their shape for 6 days (Fig. 4E). However, this only happened in a subset of cross-shaped tissues with 2 mm-long branches and printed with 200,000 cells per mm^2^ of pattern. Crosses with lower cell density could experience structural failure, with one (Fig. 4F) or two (Fig. 4G) branches breaking after a couple of days. Given that myoblasts tend to align to fuse while simultaneously generating uniform forces in the same direction, it is likely that increased contractility in two different directions induced shear stress, eventually leading to tissue rupture. It suggests an intrinsic tendency of the tissues to reorganize into a more physiologically relevant uniaxial geometry.

Increasing the initial cell density from 100,000 to 200,000 cells per mm^2^ improved tissue cohesion and reduced the likelihood of breakage. However, it also makes the tissue more likely to detach from the needles, as it would be expected that the adhesion area remains constant despite the increased cellular tension of thicker tissues. To mitigate this issue, additional NP3-labeled C2C12 cells were pipetted towards the magnetized needles, forming a reinforcing shell around each clamp. This approach effectively strengthened both the tissue structure and its attachment points, enabling the stable culture of clip-on cross-shaped tissues without breaking for at least one week, with a reproducibility rate exceeding 80%, even for crosses with 4 mm-long branches (Fig. 4H-I).

With the successful stabilization of the cross geometry, it became possible to examine tissue maturation. Although the overall cross shape was maintained over time, the branches progressively became thinner (Fig. 4J, Supplementary Movie 3) following an exponential decay with a time constant of approximately two days (Fig. 4K). While the central region exhibited no clear anisotropy (Fig. 4L), cells positioned further from the center displayed increasing alignment along the direction of the branches (Fig. 4M). Since alignment is a key factor in myogenesis, differentiation in the branches was further evidenced by fused multinucleated myotubes, and validated using myogenin staining. Myogenin, a key transcription factor in skeletal muscle differentiation, was found to be more expressed in the branches of the tissues than their centers (Fig. 4N), with signal intensity plateauing at about 50% above central levels (Fig. 4O and Supplementary Fig. 5).

Interestingly, in tissues where one (Fig. 4P) or two (Fig. 4Q) branches had broken, myogenin was overexpressed in the center. This suggests that the mechanical stress in this region was significant, and when combined with tissue compaction due to the retraction of the broken branch(es), may have favored muscle differentiation. Additionally, cell alignment and especially myotube organization at the branches’ junction in these damaged tissues indicate an apparent attempt to reconnect the remaining branches.

This observation leads to an important insight: while we have demonstrated that non-physiological cross-shaped geometries can be imposed and stabilized using magnetic bioprinting, highlighting the versatility of this approach in magnetic tissue engineering, the phenomenon of a single central branch forming suggests an intrinsic property of the cells. Specifically, as myoblast alignment in a single direction is crucial for myogenesis, the cross-shaped tissues appear to be attempting to reorganize into more physiologically relevant fascicle-like structures. This natural tendency towards alignment prompted us to shift focus towards more uniaxial tissue geometries, which we believe will allow for a deeper understanding of the properties and behavior of magnetically printed tissues.

### Magnetic trapping and stretching enable mechanical characterization of printed tissues

The chosen wrench-shaped tissue [53], designed with a uniaxial central fiber and two clamps, also exhibited progressive spherification over the course of a week (Fig. 5A and Supplementary Fig. 6), curling up on itself. However, this geometry not only better replicates *in vivo* tissue organization but also facilitated the tissue’s magnetic trapping between two needles (Fig. 5B), within one of the compartments of the Ecoflex chip [53]. More importantly, this shape allowed for a more thorough exploration of the properties of the magnetically printed tissues, which had previously only been cultured under static geometrical constraints. To introduce dynamic stresses, which are essential *in vivo*, and evaluate how magnetic printed tissues would respond, the chips were clamped onto an automated motorized stretcher (Fig. 5C) after an overnight incubation of the tissues on the collagen-coated needles. Stretching the chips resulted in direct mechanical deformation of the tissues without detachment in most cases, as adhesion to the needles was established during incubation. This setup provided a valuable platform for assessing the mechanical properties of printed tissues. As observed with cross-shaped tissues, printed tissues can rupture, and evaluating their resistance to rupture offers insight into their cohesion and internal organization. This is particularly relevant given that wrench-shaped tissues resemble tensile test specimens commonly used in material testing.

**Figure 5.**
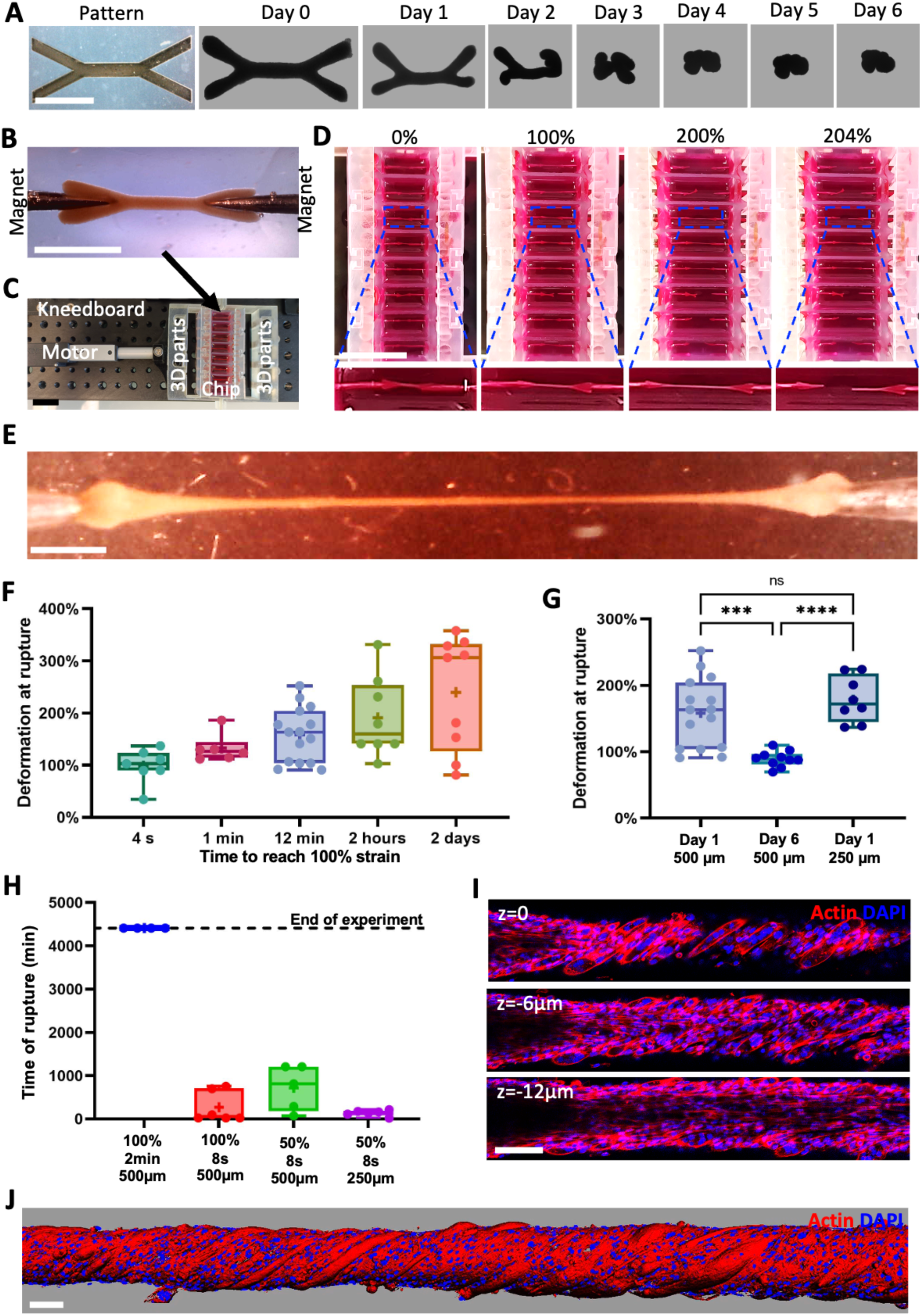
Mechanical resistance of magnetically printed tissues under stretching. (A) Brightfield images showing the evolution of free-floating tissues printed with wrench-shaped patterns (100,000 cells per mm^2^ of pattern) from day 0 to day 6. (B) Magnetically trapped clip- on wrench-shaped tissue on day 1. (Scale bar = 2 mm) (C) Automated stretching system and (D) progressive stretching of a chip until tissue rupture. (Scale bar = 2 cm) (E) Tissue stretched at a rate of 100% strain over 2 days on day 6, reaching four times its original length. (Scale bar = 1 mm). (F-G) Deformation at rupture of (F) 1-day-old C2C12 tissues printed with 500 µm-wide patterns, stretched at rates of 100% strain over 4 s, 1 min, 12 min, 2 hours, and 2 days, and (G) comparison of rupture deformation for tissues stretched at a rate of 100% strain over 12 min on day 1 versus day 6, printed either with 500 µm- or 250 µm-wide patterns. (H) Time of rupture for tissues subjected to cyclic stretching from day 1 with 100% strain amplitude and 2-minute cycles (500 µm-wide patterns), 100% strain amplitude and 8-second cycles (500 µm-wide patterns), and 50% strain amplitude within 8-second cycles (500 µm- and 250 µm-wide patterns). (I) Confocal imaging of multiple z-slices and (J) 3D reconstruction of a tissue stretched from day 1 to day 4 with 100% strain amplitude 2-minute cycles. Actin is shown in red and nuclei in blue. (Scale bar = 100 µm)

One of the first parameters examined was the impact of stretching speed on tissue rupture. The automated stretching system allowed tissues to be elongated at different speeds, ranging from 100% deformation over a few seconds to gradual deformation over several days, while monitoring at what point each tissue would rupture (Fig. 5D, Supplementary Movie 4). Tissues stretched at a speed of 100% over two days reached over four times their original length (Fig. 5E) before rupture, whereas those subjected to rapid stretching could only extend to approximately twice their initial length (Fig. 5F). Although such strains exceed physiological levels, these results still indicate that allowing cells to adapt during stretching with a lower stretching speed enables them to withstand significantly higher strains. However, when tissues were stretched at an intermediate speed (100% over 12 minutes), those stretched only after 6 days of differentiation could barely be stretched more than 100%, whereas tissues stretched on day 1 could reach more than 200% strain (Fig. 5G). To confirm that this was not due to the tissues becoming thinner, from 358 ± 38 µm on day 1 to 203 ± 53 µm on day 6, rupture experiments were conducted on day 1 using tissues printed from patterns with a 250 µm-wide central fiber, which was half the width of the control (Supplementary Fig. 7). These tissues, with an average width of 238 ± 30 µm on day 1, exhibited similar rupture behavior to the control (Fig. 5G). This suggests that the increased brittleness of tissues on day 6 is a result of their maturation process and tissue reorganization, making them less capable of withstanding deformation unless they have been progressively stretched over time.

Automating the stretching system also enabled the application of more complex stretching protocols, like cyclic stretching. To complement studies focusing on physiological ranges of deformation [54–59], we applied cyclic stretching with larger amplitudes (100% and 50%) to further test the mechanical properties of the printed tissues over four days. Interestingly, tissues stretched to 100% amplitude within 2-minute cycles (ramping from 0% to 100% in 1 minute, then returning to 0% in 1 minute) did not rupture over the four-day experiment. In contrast, tissues exposed to 100% strain in 8-second cycles broke within a few hours at most (Fig. 5H). Reducing the amplitude to 50% more than doubled the time before rupture, indicating that while strain amplitude does play an important role, cycle duration is more critical. Additionally, tissues printed with 250 µm-wide central fibers ruptured five times faster than those with 500 µm-wide fibers, even though tissue width did not influence the maximum deformation under linear stretching. This suggests that rupture under cyclic stretching may result from material fatigue rather than exceeding the limit of deformation. Notably, tissues that did not rupture under 100% strain within 1-minute cycles displayed a distinct internal organization, with cells aligning at a 45° angle relative to the overall tissue structure (Fig. 5I-J). Given that they were alternately subjected to tensile forces (which align cells along the stretching axis) and compressive forces (which promotes perpendicular alignment), the 45° orientation likely represents an optimal configuration that minimizes overall stress.

### Magnetic cell segregation enables spatial organization in co-culture tissues

After characterizing the geometrical and mechanical properties of tissues composed solely of C2C12 myoblasts, we introduced a second cell type - NiH3T3 fibroblasts - during the bioprinting process. Myoblasts and fibroblasts were respectively mCherry and GFP Lifeact transfected to distinguish them. Additionally, the cells were magnetically labeled using two distinct nanoparticle incubation protocols: NP3 labeling or NP1 labeling (a single 30-minute nanoparticle incubation the day before bioprinting). With a single incubation, myoblasts internalized slightly more nanoparticles (8 pg per cell) compared to fibroblasts (6 pg per cell) (Fig. 6A). As expected, this difference became more pronounced after two additional incubations, with myoblasts reaching an average of 20 pg per cell, against 11 pg for fibroblasts.

**Figure 6.**
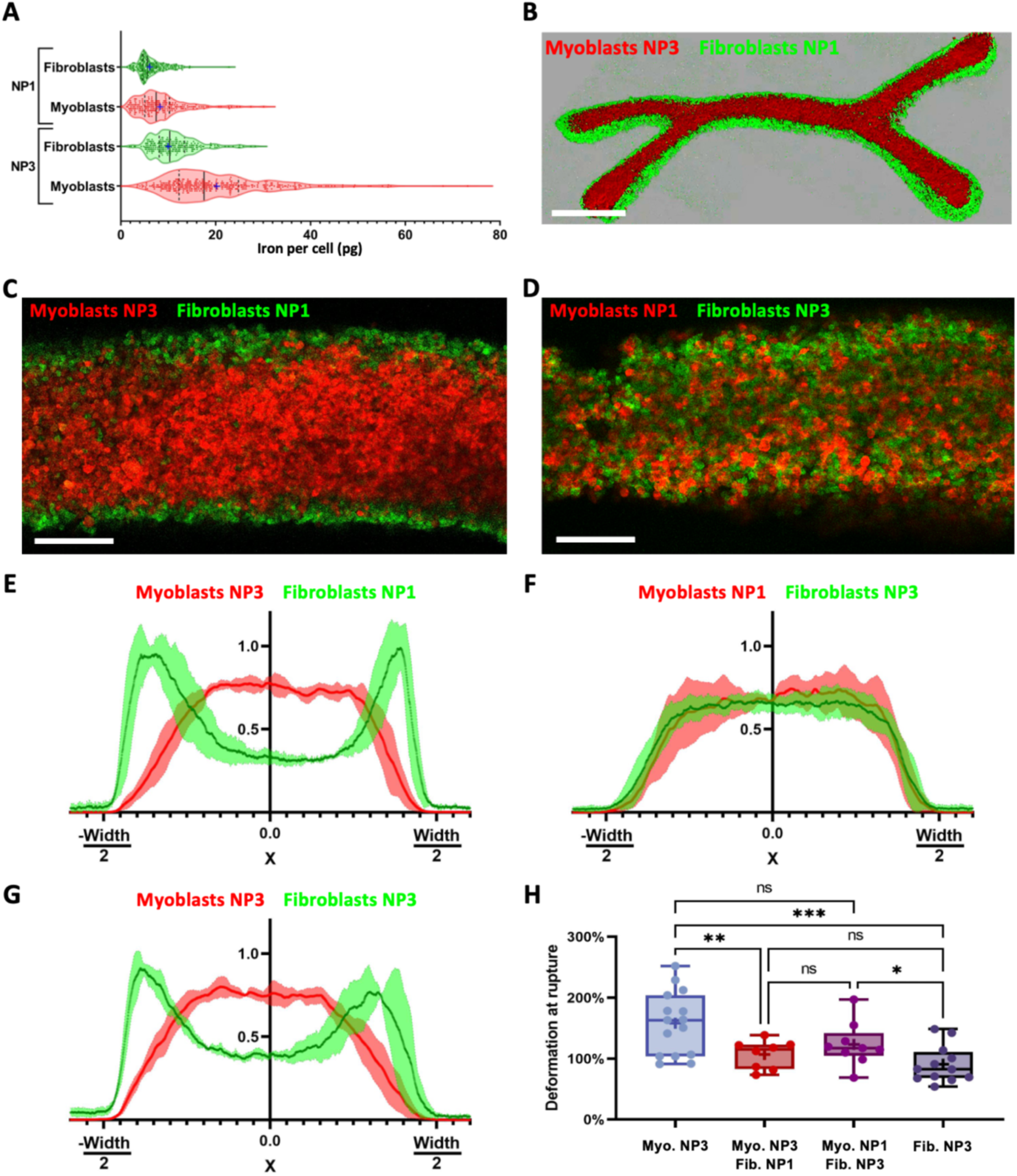
Magnetic segregation in bioprinted tissues composed of two cell types. (A) Distributions of iron mass internalized by C2C12 myoblasts and NiH3T3 fibroblasts following NP1 or NP3 labeling. (B) 3D reconstruction from confocal imaging of a tissue printed with 50% NP3-labeled myoblasts cells and 50% NP1-labeled fibroblasts. (Scale bar = 1 mm) (C-D) Confocal z-slices of tissues printed with (C) 50% NP3-labeled myoblasts and 50% NP1-labeled fibroblasts, and (D) 50% NP1-labeled myoblasts and 50% NP3-labeled fibroblasts. (Scale bar = 200 µm) (E-G) Normalized fluorescence intensity profiles of mCherry-Lifeact myoblasts and GFP-Lifeact fibroblasts along the width of the central fiber in tissues composed of (E) 50% NP3-labeled myoblasts and 50% NP1-labeled fibroblasts (n=4), (F) 50% NP1-labeled myoblasts and 50% NP3-labeled fibroblasts (n=5), and (G) 50% NP3-labeled myoblasts and 50% NP3-labeled fibroblasts (n=4). (H) Deformation at rupture on day 1 for tissues stretched at a strain rate of 100% in 12 minutes, comparing tissues composed of NP3-labeled myoblasts, NP3-labeled fibroblasts, 50% NP3-labeled myoblasts and 50% NP1-labeled fibroblasts, and 50% NP1-labeled myoblasts and 50% NP3-labeled fibroblasts.

When NP1-labeled fibroblasts and NP3-labeled myoblasts were mixed in equal proportion before bioprinting, their difference in magnetization led to their spatial segregation within the printed tissue. The more magnetic myoblasts accumulated primarily at the center of the structure, while the less magnetic fibroblasts localized at the periphery (Fig. 6B-C). This phenomenon likely occurs because the more magnetic myoblasts move faster and reach the whole patterned area first, while the less magnetic fibroblasts accumulate afterwards where the magnetic field gradient is strongest - along the edges of the pattern. In contrast, when the labeling was reversed (NP3-labeled fibroblasts and NP1-labeled myoblasts), the distributions of iron internalization of the two cell types were very similar, leading to a random distribution in the tissues after magnetic bioprinting (Fig. 6D). This segregation effect was further confirmed by fluorescence intensity profiles across tissue width which showed a distinct central localization of myoblasts when they were more magnetic and a uniform distribution when their magnetization was lower (Fig. 6E-F). Intermediate levels of segregation were observed when different labeling combinations were tested, depending on the relative difference in magnetization between the two cell types (Fig. 6G and Supplementary Fig. 8).

Rupture tests were performed on tissues composed solely of NP3-labeled fibroblasts, at a strain rate of 100% in 12 min. These tissues broke at much lower deformation compared to myoblast-only tissues, prompting an investigation into the mechanical behavior of mixed-cell tissues. Interestingly, tissues where the two cell types were magnetically segregated exhibited properties closer to those of the fibroblast-only condition, whereas tissues with a random distribution were more similar to myoblasts-only tissues (Fig. 6H). One possible explanation is that when the tissue is segregated, the weaker fibroblast region dictates the moment of rupture, as it breaks first, leading to overall failure. However, when the two cell types are homogeneously mixed, the more deformation-resistant myoblasts may help compensate for the lower mechanical strength of fibroblasts, delaying tissue rupture.

## Discussion

In this study, we demonstrate how magnetic-based approaches, particularly magnetic bioprinting, can be used to control both the macroscopic shape and microscopic cellular architecture of engineered tissues. Reproducing specific macroscopic geometries is a key objective in 3D tissue engineering. Many approaches focus on a single predefined shape for a specific application, such as moulding collagen tubes to mimic tubular structures [12] or muscle fascicles [13,17,18], or stamping intestinal-like grooves onto matrices [11]. Bioprinting [20,21,23], relying on ink extrusion or photopolymerization, increases the versatility and complexity of the designs that can be created, sometimes closely resembling organs [25]. These techniques almost systematically depend on scaffolds, either preloaded with cells – subjecting them to stress during gel extrusion - or seeded post-fabrication, requiring cellular colonization over time [25,60]. Magnetic bioprinting differs by being scaffold-free, allowing unrestricted cell density. While previous studies on magnetic bioprinting have primarily achieved simple geometries [31,46–48] such as rods, rings or spheroids, the method introduced here enables a level of 2D complexity comparable to conventional bioprinting techniques. Additionally, tissue thickness can be adjusted to provide a degree of 3D control, though not as advanced as extrusion-based techniques. Importantly, this approach inherently imparts magnetic properties to the printed tissues, allowing them to be actuated by magnetic forces, as evidenced by their ability to clamp onto magnetic needles and undergo stretch-induced stimulation. This new set of magnetic tools resembles 4D bioprinting [61], where scaffolds can evolve in response to stimuli, sometimes even magnetic [60]. But our approach can bypass the matrix and directly control the cells.

Controlling the macroscopic shape of the tissue also influenced internal cellular organization [3]. The clearest example is the clip-on cross tissues, where cells aligned along the four branches but remained unorganized in the center. This pattern likely arises from the mechanical forces within the tissue, as cells tend to minimize deformation energy [62,63]. With one end constrained by a needle and the other connected to the center, each branch experiences traction forces that align cells along its axis, while the center, subjected to multidirectional forces, lacks a dominant orientation. This also creates shear stresses, making the center a structural weak point if the tissue is too thin. In contrast, tissues embedded in collagen were initially constrained on every external surface by the surrounding matrix. Detaching from this matrix allowed the branches to compact and develop compression forces, which led to cell perpendicular alignment. These conditions under static constraints suggest that cells in 3D tissues align based on boundary constraints, consistent with previous observations [59], but now demonstrated in a more complex geometry. Under dynamic constraints, stretching wrench shaped tissues caused C2C12 cells to adopt an oblique 45° orientation relative to the tissue and stretching direction. While traction induces parallel alignment [31] and compression leads to perpendicular alignment, the 45° orientation may represent an intermediate state that minimizes overall stress. Most studies on cyclic stretching show that cells can align perpendicularly to the stretching direction in 2D [57–59], while in 3D, alignment varies: cells may be perpendicular [54,55], parallel [56], or even position-dependent - parallel in the core and perpendicular at the periphery [54]. In our constructs, outer cells twisted around the tissue, adopting an oblique orientation. This 45° orientation is uncommon in 3D tissues and may result from the high stretching amplitude (100%), whereas most studies do not exceed 20% [59]. Another key distinction is the absence of a matrix, which in traditional scaffolds provides guidance cues that reinforce alignment. Matrix remodeling can even amplify alignment over time, leading to a self-reinforcing organization [64]. By observing these diverse cellular orientations in scaffold-free tissues, we gain insight into how structural and mechanical cues - purely from cell-cell interactions - govern cell alignment. From a tissue engineering perspective, this study showcases strategies to actively shape tissue architecture and control cellular orientation in multiple directions.

Beyond cell orientation, cellular organization within tissues could also be controlled through magnetic segregation of multiple cell types. In microfluidic chips, co-culture is typically achieved by isolating different cell types in separated compartments, whereas precisely arranging multiple cell types within a single tissue remains challenging. One effective method involves extruding multiple gels, each loaded with a different cell type [65], offering versatility but adding complexity to the bioprinting process. In contrast, co-culture does not complicate magnetic bioprinting. While less versatile than extrusion-based approaches, it enables a simple and scalable way to achieve core-shell tissue organization in various geometries, which holds potential for tissue engineering. A previous study used matrix-guided moulding to create endothelial vessels surrounded by muscle cells, but only in a uniaxial configuration [18]. With the increased geometric flexibility provided by magnetic bioprinting, more complex tissues could be engineered, such as tubular networks embedded within another cell type, or multilayered structures mimicking smooth muscle layers in the intestines for instance.

Controlling both the geometry and internal organization of tissues is not only interesting for tissue engineering but also provides insight into how these parameters influence tissue mechanical properties, particularly fracture mechanics. Proper resistance to rupture is essential during development, as tissues experience a variety of stresses, and can serve to distinguish healthy from pathological tissues. It is generally assumed that under mechanical stress, a tissue stretches until the deformation exceeds the adhesion forces between cells or matrix components, at which point an initial defect forms and propagates through cell junctions, eventually to the entire tissue [66]. This failure mechanism is well documented in cell monolayers and 2D systems [67,68]. However, fewer studies have investigated it at the level of a 3D tissue. In 3D, some studies on the separation of cellular aggregates [69] have examined tissue fracture, yet primarily focusing on the cellular [70] and adhesion molecule [71] scales. Magnetic printed tissues offer a new tool to study tissue cohesion and resistance, notably thanks to their adaptable design. Our findings demonstrate that tissues of different widths (500 µm and 250 µm) showed similar rupture distributions, confirming that deformation at failure under linear stretching is a material property independent of geometry, as previously observed in epithelial monolayers [67]. Moreover, we observed a decrease in deformability at higher strain rates, consistent with prior studies. However, we extended our analysis to much lower strain rates, including experiments lasting more than an hour, and even up to a week. At these timescales, cellular rearrangements occur, making it difficult to assess purely physical material properties, yet the trend remained: lower strain rates corresponded to higher rupture deformation. Interestingly, slow stretching appeared to condition the tissue, making it more stretchable than tissues subjected to rapid deformation after six days. Beyond strain rate effects, fracture mechanics in tissues composed of two cell types - especially when spatially segregated - has not been investigated to the best of our knowledge. The difference in rupture behavior between segregated and homogeneously mixed tissues observed in this study can be explained by a crack propagation model [66], which assumes that a crack appears when the energy needed to deform a cell exceeds the energy needed to break cell adhesion. In randomly distributed co-cultures, the more deformable myoblasts can compensate for the higher brittleness of fibroblasts, resulting in an overall intermediate mechanical response. However, in segregated tissues, fibroblast-rich regions must stretch the same extent as the rest of the tissue. Since fibroblasts are less deformable, cracks form earlier in these weaker regions and propagate through the tissue, ultimately leading to premature failure.

Regarding cyclical stretching, it is typically used in mechanostranduction studies [31–35] to promote cell differentiation. In this study, it also provided insights into the mechanics of magnetic bioprinted tissues. Indeed, despite deforming the tissues by 100%, a strain rate of 100%/min was low enough to allow cyclic stretching over multiple days, whereas a rate of 25%/s was not sustainable. This supports the idea that, unlike inert materials, tissues can repair themselves, with cells capable of re-stablishing adhesion given sufficient time [72]. For example, in *Drosophila* tissue subjected to pulsatile forces [73] myosin-mediated adhesion repair occurs within 400 seconds. Despite this regenerative ability, living tissues remain susceptible to mechanical fatigue like conventional materials[72]. Repeated sub-failure loads lead to the accumulation of micro-defects, which can propagate and cause rupture. Higher strain amplitudes increase the likelihood of such defects appearing [74], and unlike linear stretching, tissue geometry influences fatigue-induced failure - thinner constructs are more prone to breakage under cyclic loading [75].

Together, these findings highlight the need to consider both mechanical and biological factors when studying the properties of living tissues. By controlling tissue architecture at both the macroscopic and microscopic levels, as well as the mechanical strains applied, the magnetic-based approaches in this study demonstrate potential not only for tissue engineering but also for advancing our understanding of the biomechanical principles governing living tissues.

## III. Methods

### Cell culture

C2C12 cells (ATCC) were cultured in growth medium, DMEM (Gibco) with 10% fetal bovine serum (Dutcher) and 1% Penicillin-Streptomycin (Thermofisher).

NiH3T3 fibroblasts (ATCC) were cultured in growth medium, where fetal bovine serum was replaced by calf serum (ATCC 61965-026).

Both cell types were passaged once confluency reached between 50 and 80%, without exceeding passage 5 for C2C12 and 10 for NiH3T3.

### Cell Transfection

Cell transfection was performed using the PEI MAX reagent (Polysciences) according to the manufacturer’s protocol. The pLVX-LifeAct-mScarlet plasmid and the pLVX LifeAct-eGFP-P2A-puro plasmid (constructed from pLVX puro vector Clontech Catalog No. 632164) were used for viral transfection. The pLVX-LifeAct-mScarlet plasmid was employed to infect C2C12 cells, generating the C2C12 LifeAct-mScarlet stable cell line. Similarly, the pLVX-LifeAct-eGFP-P2A-puro plasmid was used to infect NiH3T3 cells, resulting in the establishment of the NiH3T3 LifeAct-eGFP stable cell line.

### Magnetic Labeling

Maghemite nanoparticles, synthesized via the iron salts co-precipitation method and coated with citrate, were used to label the cells [76].

Labeling solution was prepared by mixing the nanoparticles at a concentration of 2 mM of iron and 2 mM citrate (to prevent nanoparticle aggregation) into RPMI medium (Gibco). For NP1 labeling, the cell culture medium was replaced with this nanoparticle solution and incubated for 30 minutes the day before the bioprinting procedure. For NP3 labeling, cells were incubated with the nanoparticle solution once daily for three consecutive days prior to bioprinting, as described previously [31]. These incubation protocols facilitated nanoparticle efficient internalization via endocytosis.

### NiFe pattern fabrication

Magnetron sputtering (Plasmionique) was used to deposit a conductive layer of 10 nm of titanium and 100 nm of copper on a 70 x 50 mm^2^ Corning glass slide. TI-Prime (Microchemicals) and then 70 µm of AZ 125nXT photoresist (Microchemicals) were spincoated on the coated side of the slide. Photolithography was then performed with a chromium mask, designed with the shape of the desired patterns. This exposes the copper in these shapes, making it possible to perform NiFe (80:20) electrodeposition [31,51,52]. The copper covered substrate, acting as a cathode, and an NiFe anode (Goodfellow) were connected to a generator while being immersed in a 250g.L^-1^ NiSO_4_, 5g.L^-1^ FeSO_4_, 25g.L^-1^ boric acid, 2g.L^-1^ saccharin and 0.1g.L^-1^ sodium dodecyl sulfate bath, heated to 30 °C and magnetically stirred. Applying a current of 7 mA per mm^2^ of patterns led to NiFe deposition in the holes with the exposed copper. In these moulds, NiFe patterns would grow by approximately 3 µm.h^-1^, until patterns would be at least 50 µm thick. The substrate would then be immersed in TechniStrip P1316 (Technic) at 70 °C to remove the copper and resist, exposing the freshly electrodeposited 50 µm thick NiFe patterns in the same shape as the original patterns on the chromium mask. The substrate was finally spincoated with 100 µm of PDMS (Sylgard 184, 1:10 curing agent). Multiple patterns were made on the same glass substrate. They were positioned to be grouped within 16 mm wide circles, the size of the wells for the next bioprinting step.

### Magnetic bioprinting process

Wells were made by plasma-bonding 1 cm-high hollow PDMS (Sylgard 184, 1:10 curing agent) disks with outer and inner diameters of 20 mm and 16 mm respectively, to a 150 µm thick 22 x 22 mm^2^ VWR coverslip. After treating them with anti-adherence solution (StemCell Technologies) for 1 hour, they were placed directly on top of NiFe patterns. 500µL of a suspension of previously magnetically labeled cells, detached with TrypLE Express 1X (Gibco) was pippeted into each well. The number of cells in this volume was adapted to the total area of the NiFe patterns beneath the wells, aiming for either 10^5^ or 2×10^5^ cells per mm^2^ of pattern. The substrate with the patterns and the wells on top was then placed in the middle of two magnets (110.6 × 89 × 19.6 mm^3^, remanence Br = 1.35 T, Ref. Q-111-89-20-E, Supermagnete), positioned to attract each other but maintained at 3.6 cm, a distance meant to minimize the magnetic gradient [31]. After usually 3 hours the system was removed from the magnets and the magnetic printed tissues were collected.

### Tissue culture after bioprinting

Following bioprinting, C2C12 cells were cultured in differentiation medium, consisting of DMEM (Gibco) with 1% Horse Serum (Thermofisher) and 1% Penicillin-Streptomycin (Thermofisher).

To investigate whether the patterns could prevent tissue compaction, some tissues were maintained on the patterns within the magnetic field for 6 days, without changing the initial configuration from day 0. For all other conditions, tissues were collected from the wells with a 5 mL pipette just after removing them from the magnets and the patterns, 3 hours post-bioprinting. The tissues were then cultured for 6 days, either free floating in wells of non-treated VWR 48-well plates, magnetically trapped by needles in ecoflex chips or embedded within a 2 mg.mL^-1^ collagen matrix. For the latter, a layer of 100 µL of a cold collagen solution (differentiation medium with 2 mg.mL^-1^ collagen (Corning, rat tail) and pH 7 adjusted with NaOH) was put to polymerize at the bottom of a well of a 48 well plate. After placing the printed tissue on top of the polymerized collagen layer, which prevents it from sedimenting at the bottom, the supernatant was removed and a second layer of 100 µL collagen solution was added. Once polymerized after 30 min in an incubator, the tissue was fully embedded in the matrix, and the well was filled with differentiation medium.

### Chip fabrication

Ecoflex (00-31 Near Clear, Smooth-on) was poured into 3D printed moulds, designed on Autodesk inventor, with needles inserted where the final needles should be, and then placed at 75°C for 3 hours. After withdrawing the needles, the chips could be unmoulded and nickel-plated steel needles (0.6 mm wide, Bohin) were inserted back in the chips. To avoid them from rusting in cell culture medium, these final needles were covered with 3 coatings of transparent nail polish dried overnight at 120°C to evaporate all solvents. The chips for cross shaped tissues were designed as 15 mm wide square reservoirs with four needles aiming towards the center of the chips, at 2.25 mm or 4.25 mm from the center. The chips for wrench-shaped tissues were designed with 8 parallel 14 mm-long rectangular reservoirs with two aligned needles aiming towards the center and 2mm apart. Once properly placed, the needles were fixed to the chips with Loctite SI 5398 adhesive. Next, the chips, and specifically the needles, were plasma treated, exposed to Tris-HCL buffer (pH = 8.5) with 5 mg/mL dopamine hydrochloride (Thermo Scientific) for 3 hours, and then rinsed with PBS. The polydopamine treatment is meant to activate the surface of the needles for the final coating at room temperature with a PBS solution with 0.2 mg/mL collagen I for 2 hours. This coating, timed to end at the same time as the bioprinting process, was rinsed with PBS and the chips were filled with warm medium.

### Magnetic trapping process

In each chip, each needle is magnetized by a 6×6 mm N48 Neodymium magnet (Supermagnete, S-06-06-N) placed on its outward end with its axis aligned with the needle. The magnets were positioned in this specific configuration by trapping them in 3D printed slots adapted for each chip. The chip for cross shaped tissues fits into a square magnet holder with a magnet slot in the middle of each side. The magnets are arranged so that their polarity alternates, meaning that two magnets facing each other repel, creating magnetic field lines in a cross shape at the center of the chip. For wrench-shaped tissues, the chip is placed between two magnet holders, each consisting of 8 rows of magnet slots, to magnetize the two needles on each side of the 8 compartments of the chip. Within one holder, magnets are oriented identically, and magnets facing one another are oriented to attract one another, creating magnetic field lines spanning from one needle tip to the other.

For clip-on cross shaped tissues, tissues were transferred from the bioprinting well to the chip with a 5 mL pipette. As the tissue folds onto itself when aspirated, it first needs to unfold, and as soon as the magnets are placed next to the needles, each clamp of the tissue clips onto one of the four needles. For wrench shaped tissues, the magnets are already in place, and as the tissue is approached to the needles with the pipette, it magnetically clamps onto the two needles directly [53].

### 3D printed parts

Moulds for the chips and the various parts used mostly to hold magnets and stretch the chips were designed on Autodesk Inventor and then 3D printed with a Digitalwax 028J Plus® 3D printer (DWS).

### Simulations

Magnetic field simulations were done with COMSOL Multiphysics (Magnetic Field No Currents module, Licence number 6464550).

### Rupture tests

After the tissues were incubated overnight to allow adhesion to the needles, the chips were clamped onto an automated stretching system. This system was assembled using a Thorlabs breadboard, onto which motors and 3D-printed parts were fixed. One side of a chip is connected to a part fixed to the board, while the other side is connected to the motor. As the motor retracts, the chip is stretched, which increases the distance between the needles, consequently stretching the trapped tissue. The movement of the motor is controlled from a computer, via a Raspberry Pi, with a software developed to impose any type of command to the motor. The system allows up to four independent motors to operate simultaneously on one breadboard, enabling parallel stretching of multiple samples. The chips were subjected to stretching under various conditions until the tissue within each chip ruptured. During the tests, the time of rupture and the amount of stretching (when tissues were stretched linearly) were recorded. Tissues that detached from the needles before rupturing were excluded from the analysis.

### Fixation and stainings

Tissues were rinsed in PBS, immersed in 4% Paraformaldehyde in PBS (Thermofisher) for 1h at room temperature for fixation and then rinsed again with PBS. Magnetically trapped tissues were fixed as they were in their chips to preserve their shape.

Prior to staining, tissues were placed in blocking solution, 4% w/w BSA (Sigma) and 0.5% v/v Triton X-100 (Fisher) in PBS, at room temperature for 1 hour. After rinsing them with PBS they were incubated overnight at 4 °C with 1:1000 Phalloidin AlexaFluor 488 (Thermofisher) and 1:300 DAPI (Invitrogen) in blocking solution. The next day samples were rinsed with PBS.

For myogenin stainings, samples were additionally placed at room temperature in saturation solution, PBS with 0.3% (v/v) Triton X-100 and 1% (w/w) BSA. After rinsing them with PBS, samples were immersed overnight at 4°C with saturation solution with 1:200 myogenin primary antibody (mouse, DHSB, F5D). Following PBS rinsing the next day, samples were incubated at 4°C overnight with 1:750 donkey anti-mouse 488 (Thermofisher Scientific, A-21202) in saturation solution. The following day, samples were rinsed with PBS.

### Magnetophoresis for quantification of iron internalization

Magnetophoresis was used to measure the amount of iron internalized by cells following incubations with magnetic nanoparticles [77]. Magnetically labeled cells in suspension in PBS were filmed after approaching them to a magnet with a magnetic field of 0.145T and a gradient of 17 T.m^-1^. As the magnetic force attracting each cell towards the magnet should counterbalance the drag force, it can be calculated by measuring the speed and diameter of each cell. Knowing the magnetic field and gradient, the magnetic moment of each cell can then be inferred, from which the mass of iron internalized by each cell can be calculated.

### Imaging

After fixation and stainings, samples were placed on fluorodishes using tweezers and a pipette for imaging with a Leica DMi8 inverted confocal microscope employing either 10x air or 25x water objective. Live samples were imaged from below the chips or wells with a Leica DMi1 brightfield microscope for punctual images or within the incubator using a CytoSmart Lux 3 FL videomicroscopy system. A Stemi 508 Greenough Stereo Microscope with a DinoEye camera was used to image samples from above. For rupture tests, chips were imaged either by filming the entire chip during the stretching process (with the chip steadily fixed in a holder) or by taking pictures at regular intervals for stretching rates of 100% stretching over 2 hours and below. Chips that were filmed were stretched outside of the incubator for no more than 40 min, with a heating lamp to maintain a temperature of 37 °C. For longer experiments at slower speeds, the chips and stretching system were kept inside the incubator, except for medium changes and regular image capturing.

### Image analysis

Image analysis was performed on ImageJ. Images of larger objects were first reconstructed from multiple frames, either with the MosaicJ or the Stitching plugins.

Tissue shape and branch thickness were analyzed with the “Analyze particles” module.

To evaluate fluctuations of the intensity signal of myogenin in clip-on cross-shaped tissues, stacks of confocal images were summed for each channel, for each branch of the crosses a frame from the center of the tissue to the edge of the branch was cropped and the “Plot profile” module was used for the myogenin and the actin channels. For each plot, the background value was subtracted and the myogenin signal was divided by the actin signal to normalize it. The plots from all branches were averaged, resulting in a mean value of the normalized myogenin signal intensity as a function of a distance to the center. To normalize all samples together, for each of them the intensity was divided by the average value within the first 50µm from the center of the tissue.

The spatial distributions of fibroblasts and myoblasts in co-culture constructs was obtained by plotting the intensity profiles of the green and red signals along the width of the central fiber of the tissues. Both signals were then normalized in x using a python code so that the width of the tissues would be equal to 1. The minimum signal value was set to 0 and the overall signal was normalized so that the area under the curve would be equal to 1, making it easier to compare and plot the different samples together.

### Statistical analysis

Data processing was done with Microsoft Excel, and graphs were plotted with GraphPad Prism 8 (GraphPad Software, La Jolla, California). Standard deviations were used to represent error bars. Every value represented was obtained by averaging at least three independent replicates. Statistical differences were evaluated using Welch t-tests and represented with * when P value < 0.05, ** when P < 0.01, *** when P < 0.001 and **** for P < 0.0001.

## Acknowledgments

This work was funded by the French Agence Nationale de la Recherche (ANR) under grant ANR-19-CE09-0029. This work has also received the support of “Institut Pierre-Gilles de Gennes” (laboratoire d’excellence, Equipex, “Investissements d’avenir” program ANR-10-IDEX-0001-02 PSL and ANR-10-LABX-31-34), as well as funding under the MagBioPrint project grant. This work benefited from the technical contribution of the engineers of the joint service unit CNRS UAR 3750 at Institut Pierre Gilles de Gennes (IPGG) and the authors would like to thank the engineers of this unit. The authors also thank Corentin Moreau for fruitful discussions, and Sylvie Coscoy (Institut Curie) for donating transfected fibroblasts.

