## Supplementary Figures for "Shaping and probing living tissues with magnetic bioprinting"

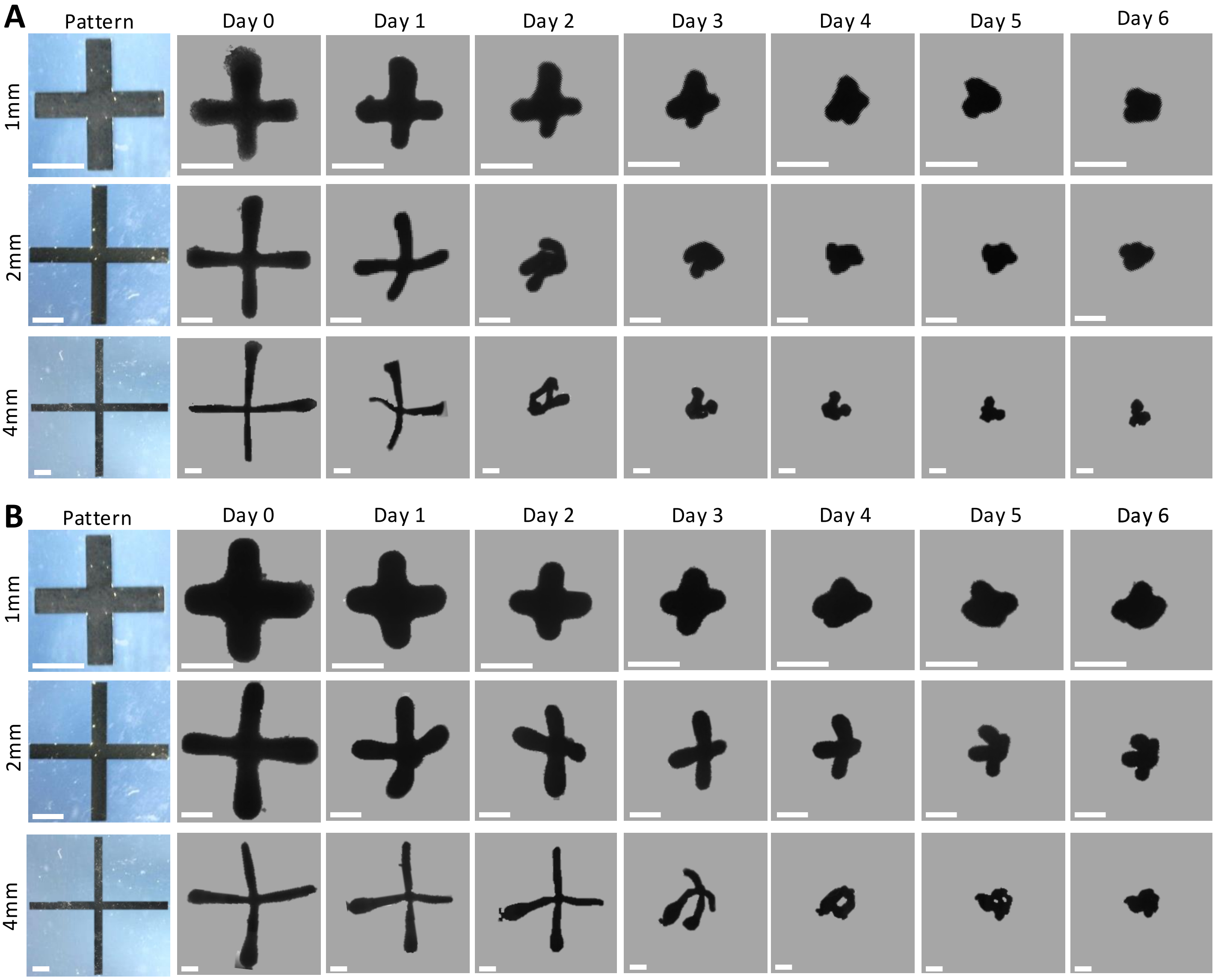


***Supplementary Figure 1.*** *Additional sets of brightfield images from day 0 to day 6 of free-floating tissues made from cross patterns with 1, 2 or 4 mm-long branches and with (A) 100000 or (B) 200000 cells per mm$^2$ of pattern. (Scale bar = 1mm)*

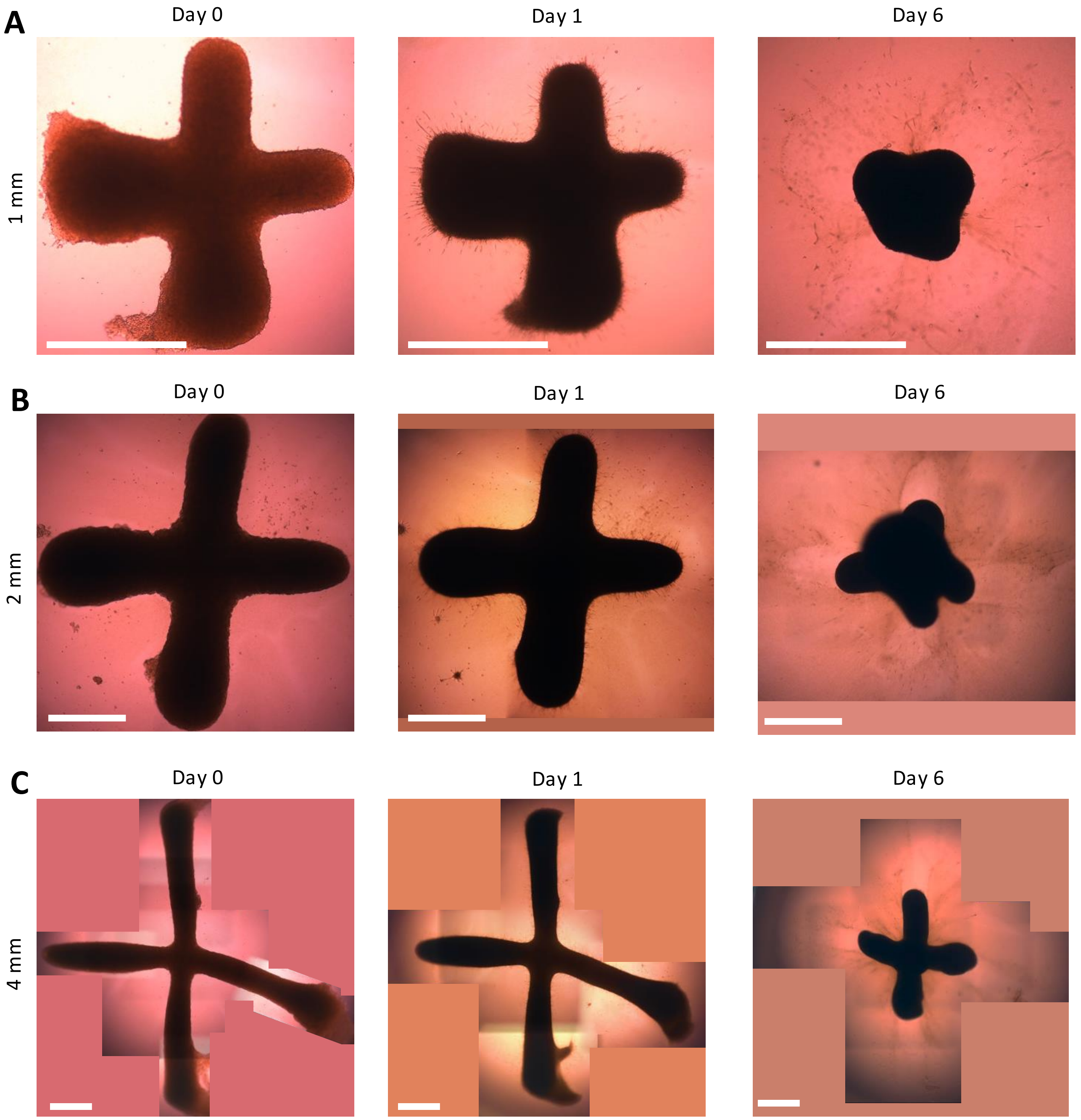


***Supplementary Figure 2.*** *Additional sets of brightfield images on days 0, 1 and 6 of tissues made from cross patterns with (A) 1, (B) 2 or (C) 4 mm-long branches and 200000 cells per mm² of pattern, embedded in collagen. (Scale bar = 1 mm) Tissues were not segmented to show the deformation of the matrix. As tissues were reconstructed from multiple frames, blank parts were filled with a uniform background color.*

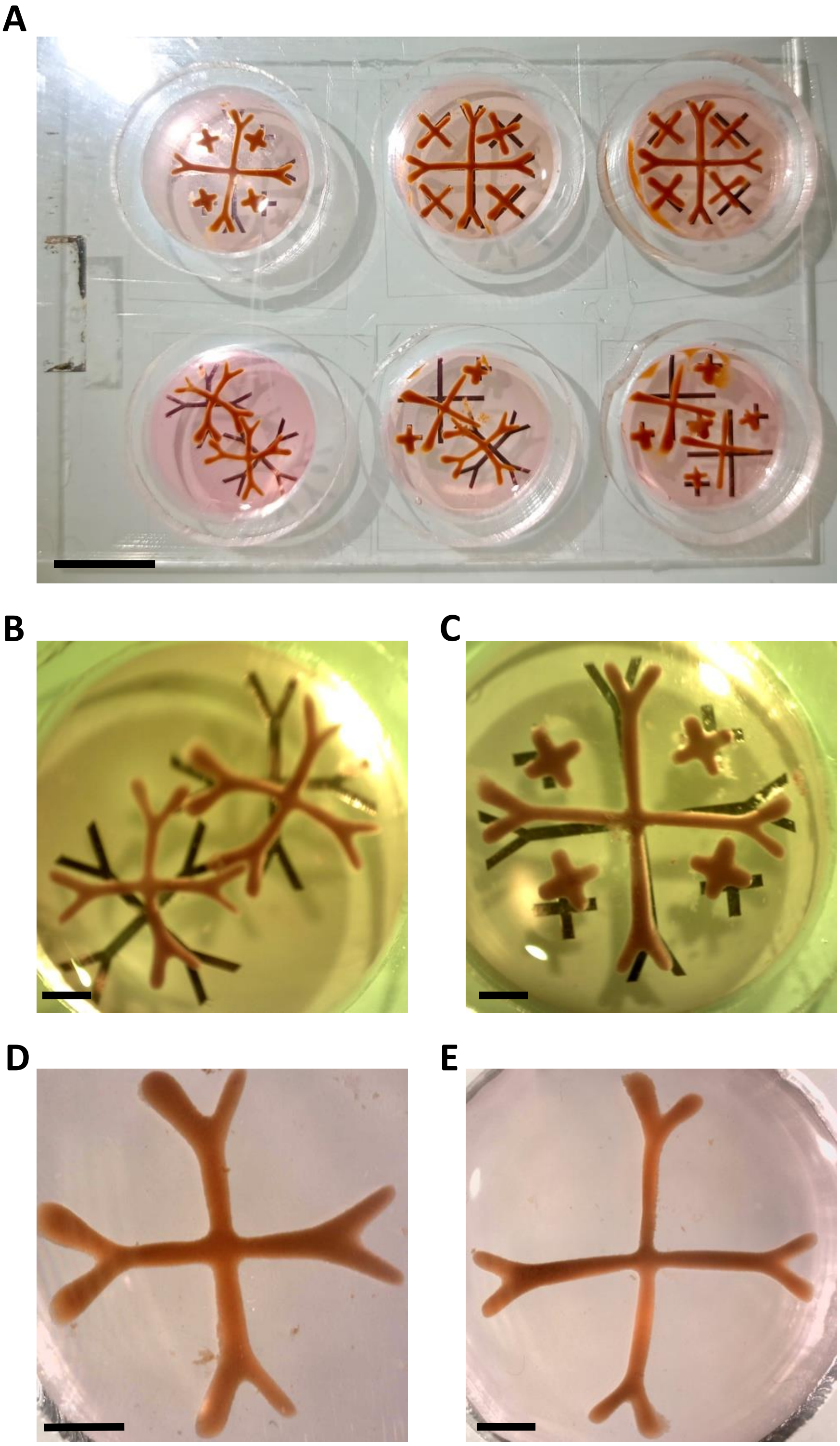

***Supplementary Figure 3.*** *(A) Image of a whole glass substrate with wells on all six sets of cross-shaped patterns, just after bioprinting (scale bar = 1cm). Wells used to make clip-on crosses with (B) 2 mm- and (C) 4 mm-long branches, with a close up on a single clip-on cross, with (D) 2 or (E) 4 mm-long branches, just after the bioprinting process (scale bar = 2 mm).*

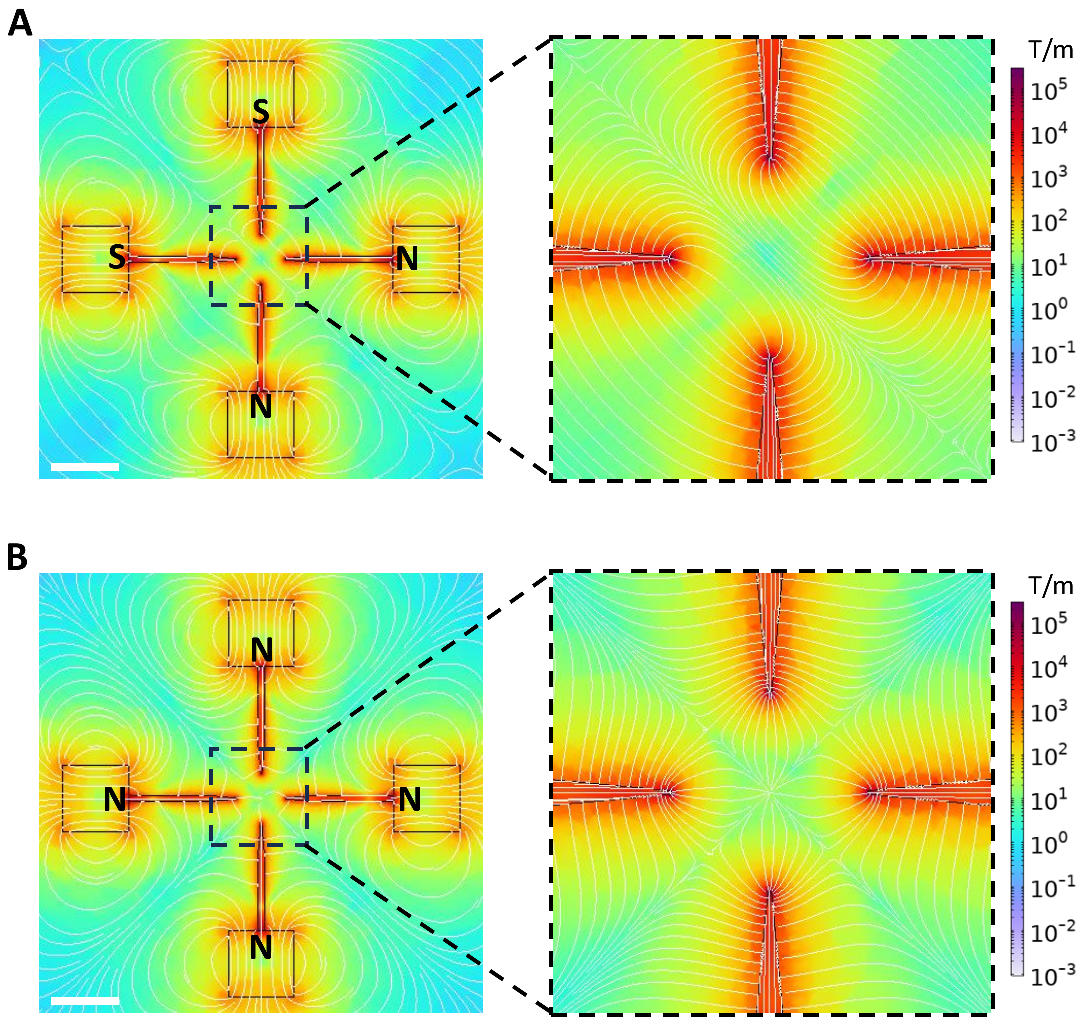


***Supplementary Figure 4.*** *Simulations of the magnetic field gradient within a chip meant to trap a cross-shaped tissue, either with (A) a North/North/South/South or (B) a North/North/North/North magnet configuration (scale bar = 1 cm).*

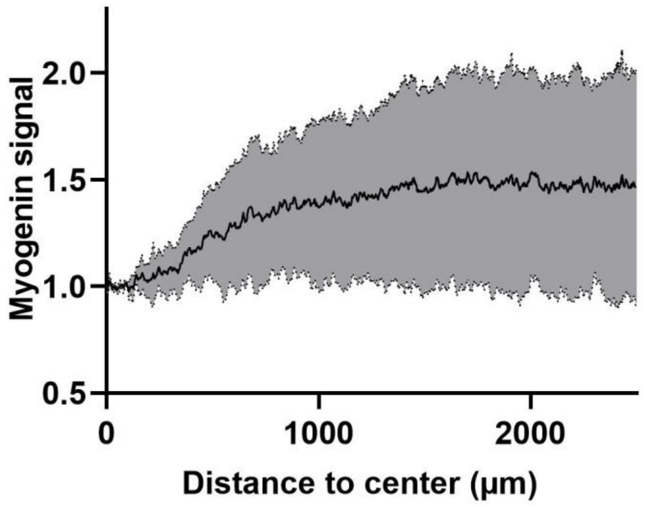


***Supplementary Figure 5.*** *Spatial distribution of myogenin within a cross-shaped tissue with 4 mm-long branches at day 6 plotted after normalization as a function of the distance from the center (n = 3)*

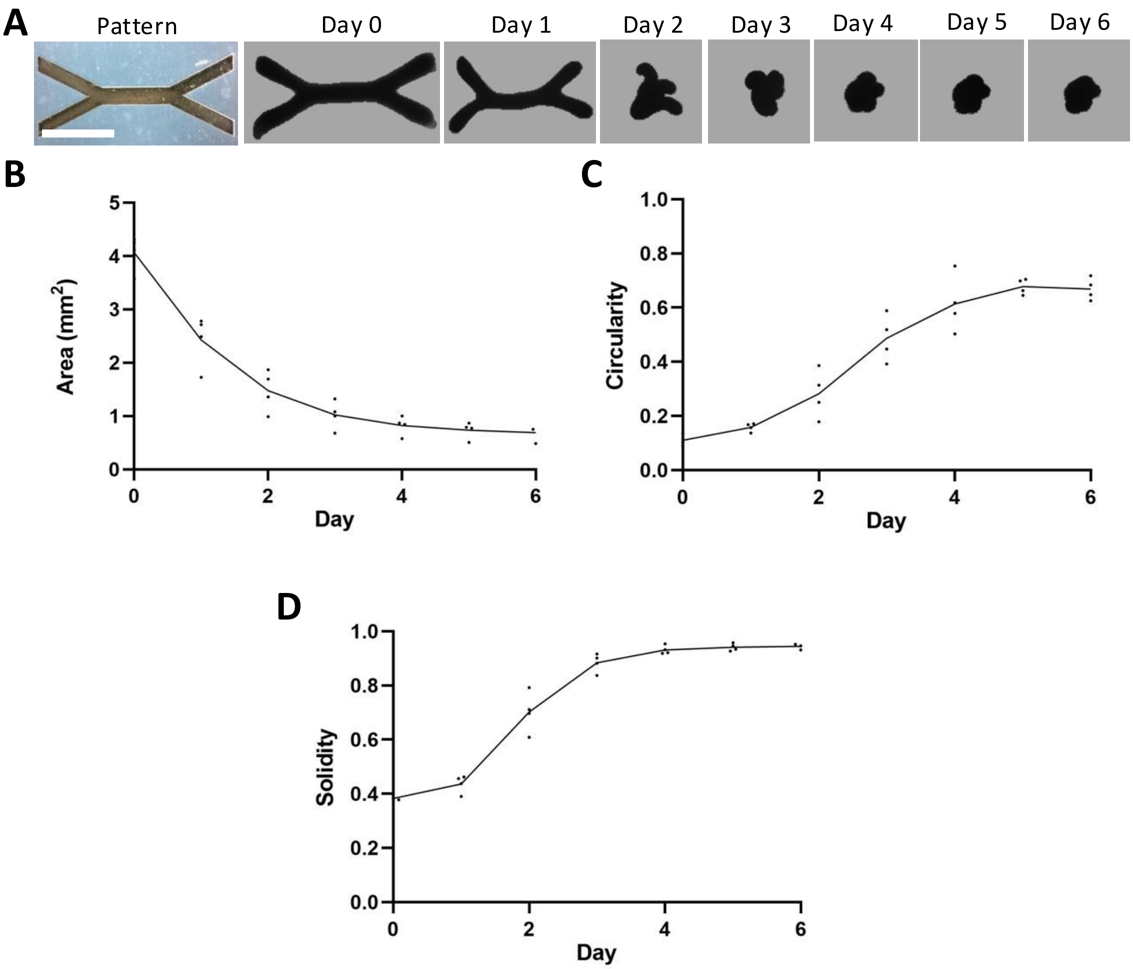


***Supplementary Figure 6.*** *(A) Additional set of brightfield images showing the evolution of a free-floating tissue printed with a wrench-shaped pattern (100,000 cells/mm²) from day 0 to day 6, and the evolution of (B) the area, (C) the circularity and (D) the solidity of wrench-shaped tissues free-floating in non adherent wells. (scale bar = 2 mm)*

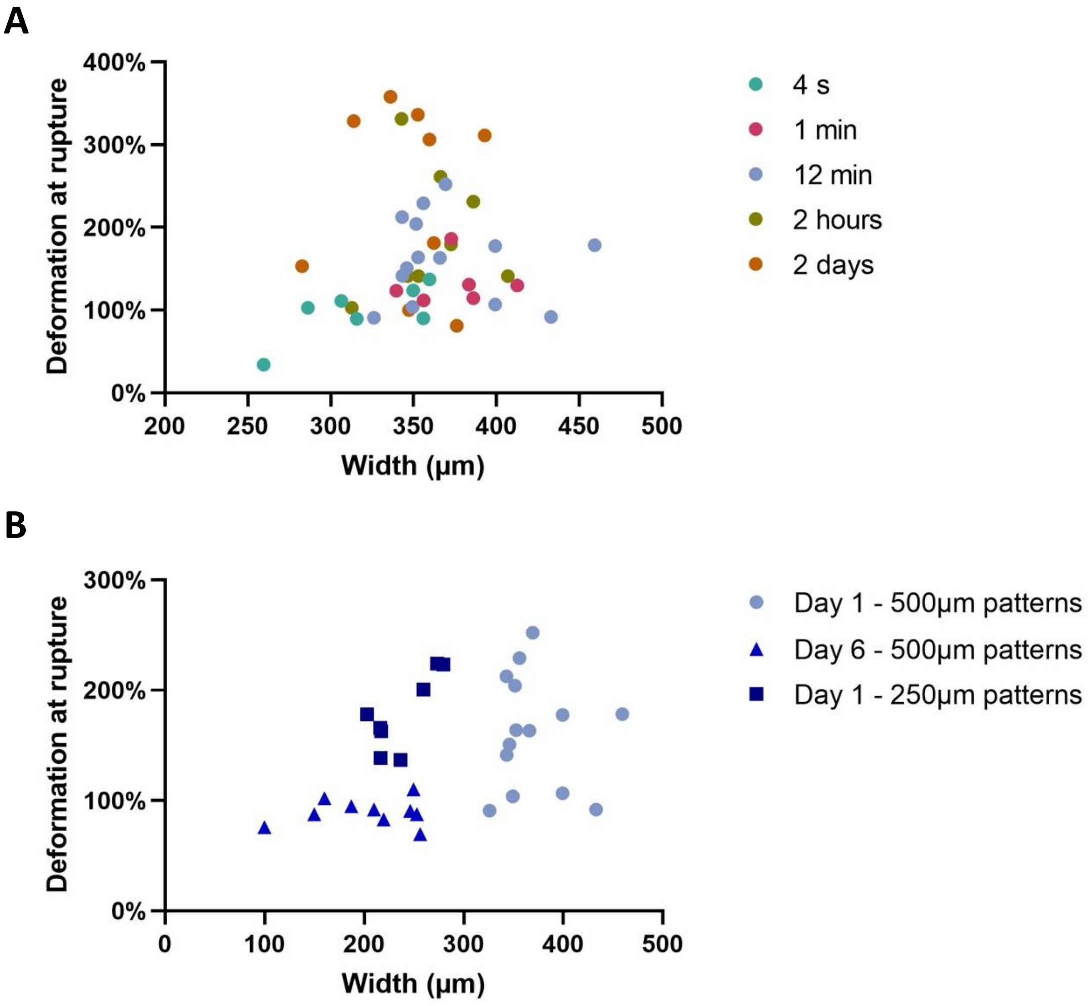


***Supplementary Figure 7.*** *(A) Maximum deformation at rupture as a function of tissue width of (F) 1-day-old C2C12 tissues printed with 500 µm-wide patterns, stretched at rates of 100% strain over 4 s, 1 min, 12 min, 2 hours, and 2 days and of (B) tissues stretched at 100% strain over 12 min on day 1 versus day 6, printed either with 500 or 250 µm-wide patterns.*

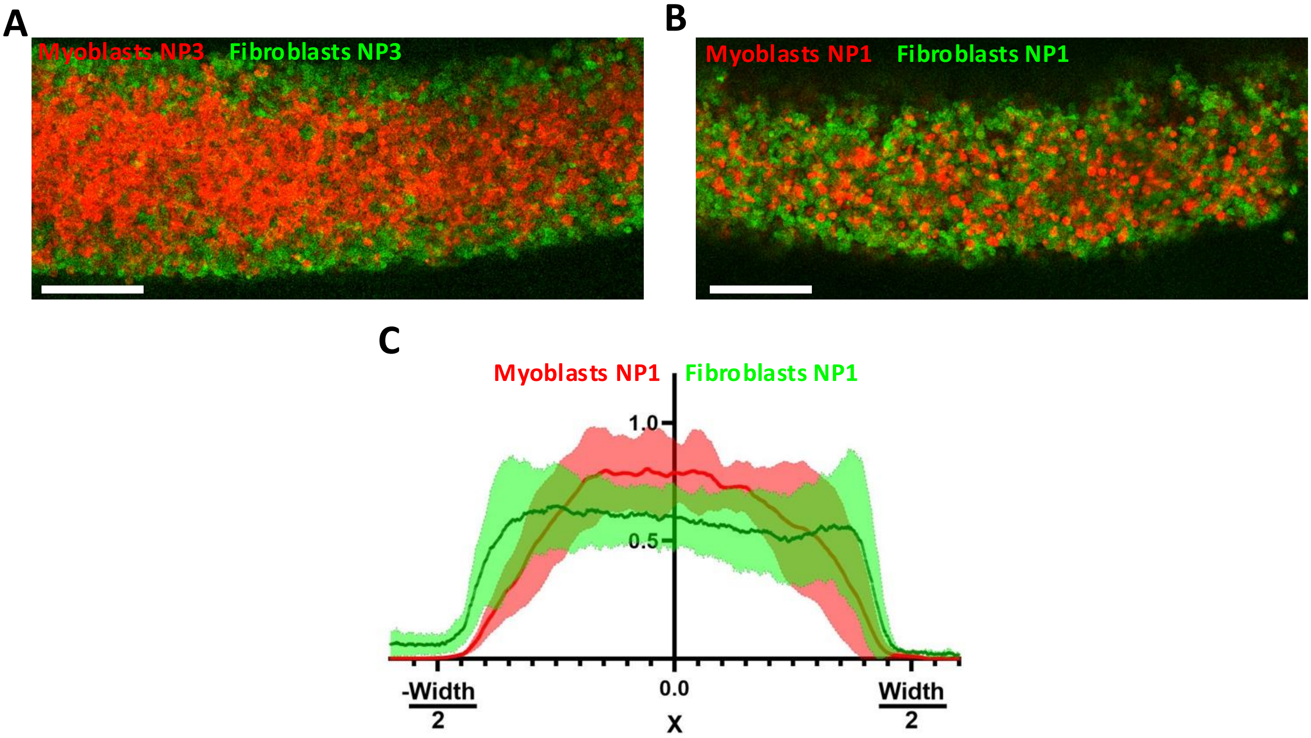


***Supplementary Figure 8.*** *(A-B) Confocal Z-slices of tissues printed with (A) 50% NP3-labeled myoblasts and 50% NP3-labeled fibroblasts, and (B) 50% NP1-labeled myoblasts and 50% NP1-labeled fibroblasts (scale bar = 200 μm). (C) Normalized fluorescence intensity profiles of mCherry-Lifeact myoblasts and GFP-Lifeact fibroblasts along the width of the central fiber in tissues composed of 50% NP1-labeled myoblasts and 50% NP1-labeled fibroblasts.*
